# Genetic association of FMRP targets with psychiatric disorders

**DOI:** 10.1101/2020.02.21.952226

**Authors:** Nicholas E Clifton, Elliott Rees, Peter A Holmans, Antonio F. Pardiñas, Janet C Harwood, Arianna Di Florio, George Kirov, James TR Walters, Michael C O’Donovan, Michael J Owen, Jeremy Hall, Andrew J Pocklington

## Abstract

Genes encoding the mRNA targets of Fragile X mental retardation protein (FMRP) are enriched for genetic association with psychiatric disorders. However, many FMRP targets possess functions that are themselves genetically associated with psychiatric disorders, including synaptic transmission and plasticity, making it unclear whether the genetic risk is truly related to binding by FMRP or is alternatively mediated by the sampling of genes better characterised by another trait or functional annotation. Using published common variant, rare coding variant and copy number variant data, we examined the relationship between FMRP binding and genetic association with schizophrenia, major depressive disorder and bipolar disorder. We then explored the partitioning of genetic association between overrepresented functional categories. High-confidence targets of FMRP were enriched for common schizophrenia risk alleles, as well as rare loss-of-function and *de novo* nonsynonymous variants in cases. Similarly, through common variation, FMRP targets were associated with major depressive disorder, and we present novel evidence of association with bipolar disorder. These relationships could not be explained by membership of other functional annotations known to be associated with psychiatric disorders, including those related to synaptic structure and function. This study reinforces the evidence that targeting by FMRP captures a subpopulation of genes enriched for genetic association with a range of psychiatric disorders, across traditional diagnostic boundaries.

## Introduction

Fragile X mental retardation protein (FMRP) binds selected mRNA species to repress their translation (1–5). In the brain, FMRP is highly, and dynamically, expressed in neurons, where it regulates the dendritic synthesis of a range of proteins (6,7), many of which are modulators of synaptic plasticity (1). The loss of FMRP function causes Fragile X syndrome (8), characterised by abnormal dendritic morphology, impaired learning and memory, autism and a high prevalence of seizures (9).

The mRNA targets of FMRP have received additional attention from psychiatric research due to their enrichment for genes harbouring risk to psychiatric disorders. A set of 842 high-confidence FMRP targets, originating from a study by Darnell *et al* in 2011 (1), have been reported to be enriched for genetic association with schizophrenia (10–16), autism (17–20) and major depressive disorder (21). In the case of schizophrenia, not only is this association robust across genome-wide association studies, but it is also mirrored across studies of multiple types of genetic mutation (common and rare) conferring risk to the disorder (10–16).

Whilst the case for the involvement of some FMRP targets in psychiatric disorders is now unequivocal, it has been noted that FMRP targets represent long, brain-expressed transcripts (22) with considerable overlap with other sets of genes enriched for genetic association with psychiatric disorders, including those encoding synaptic proteins (1,23). This has led to speculation that the association between psychiatric disorders and FMRP targets is driven not by the property of being targets of FMRP *per se*, rather that it reflects association to one or more functional sets of genes that also happen to be overrepresented in the FMRP target set (22). Furthermore, FMRP targets were defined by applying a cut-off to a probabilistic scale of FMRP binding (1), though the relationship between these binding statistics and genetic association with psychiatric disorders has not been investigated.

In the present study, we aimed to 1) establish whether the association of FMRP target genes with schizophrenia depends on binding confidence; 2) determine whether these associations can be explained by the sampling of otherwise characterised or functionally-annotated genes; and 3) demonstrate the extent to which FMRP targets are associated with risk across a range of psychiatric disorders.

## Results

### The relationship between FMRP binding confidence and enrichment for association with schizophrenia

We investigated the enrichment for common variant association with schizophrenia in bins of expressed (1) genes (N = 400 per bin) grouped by their ranking of mRNA-FMRP binding confidence. These gene set association analyses were performed using MAGMA, in which effects of gene size and SNP density are controlled for within a multiple regression model (24). Bins containing genes with greater FMRP binding confidence were more enriched for association with schizophrenia (Figure 1a), with only the top three bins being significantly associated (bin 1: corrected *P* = 2.3 × 10^−5^; bin 2: corrected *P* = 1.5 × 10^−5^; bin 3: corrected *P* = 0.030).

**Figure 1.**
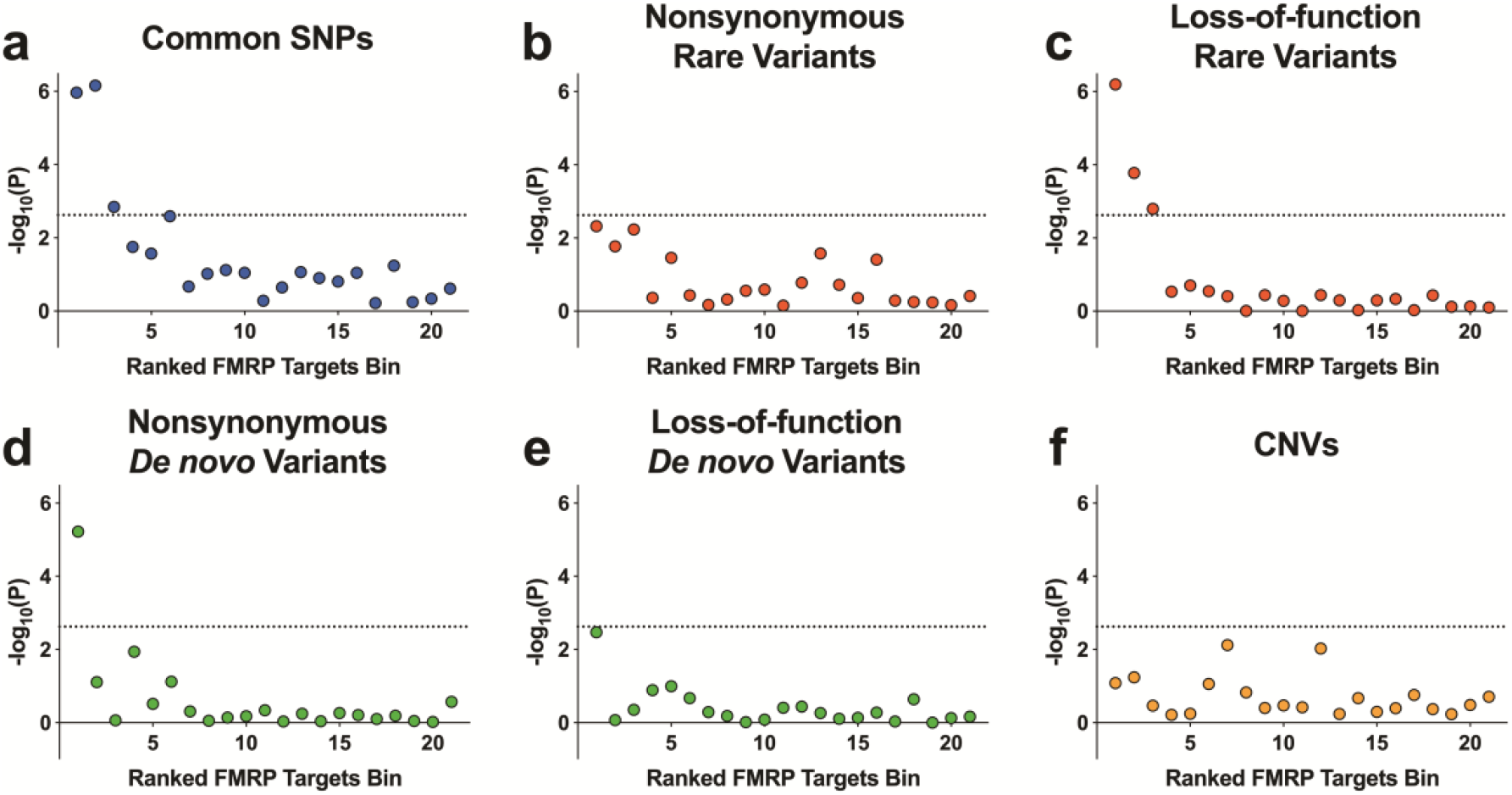
Schizophrenia association of gene sets ranked by FMRP binding confidence. All expressed genes were ranked by FMRP binding confidence and grouped into 21 bins of 400 genes. Presented are −log_10_(*P*), where the *P*-value is derived from gene set association analysis using the genetic variant type shown. CNV analyses were corrected for *P*-value inflation using random size-matched sets of expressed genes. Rare coding variants were derived from case-control exome sequencing studies of schizophrenia and defined as variants observed once in all sequenced samples and never in the non-psychiatric component of ExAC. Loss-of-function variants include nonsense, splice site and frameshift mutations. Nonsynonymous variants include loss-of-function and missense variants. Dotted lines represent a threshold for statistical significance after correction for 21 tests. SNPs, single nucleotide polymorphisms; CNVs, copy number variants.

FMRP targets have likewise been associated with schizophrenia through rare genetic variants (12–15). We used exome sequencing data to determine which bins of genes were associated with schizophrenia through rare and *de novo* coding variants. In the case-control analysis of rare loss-of-function variants, notably, the same top three bins enriched for GWAS signal were the only bins to be significantly enriched for association with schizophrenia through rare loss-of-function variants (bin 1: corrected *P* = 1.3 × 10^−5^; bin 2: corrected *P* = 0.0035; bin 3: corrected *P* = 0.034) (Figure 1c). Only the topmost bin was associated through *de novo* nonsynonymous variants (corrected *P* = 1.3 × 10^−4^) (Figure 1d).

Since risk to schizophrenia is also conferred through structural genetic variants (25–28) in the form of deletions or duplications of large sections of DNA, we investigated whether CNVs from patients with schizophrenia are enriched for genes within bins of probable FMRP targets compared to control subjects. Following logistic regression analysis, no bins surpassed the threshold for significance (Figure 1f) and the same was true if we examined deletions and duplications separately (Supplementary Figure 1).

### Refining schizophrenia association of FMRP targets through functionally defined subgroups

Many proteins translated from mRNA targets of FMRP have synaptic functions (1). In turn, substantial evidence shows that genes encoding proteins with synaptic functions are enriched for genetic association with schizophrenia(11–13,23,29,30). To further assess the importance of FMRP targeting to the association of genes with schizophrenia, we separated the 842 FMRP target genes, as determined by Darnell *et al* (1), into subgroups defined by overrepresented functional categories.

Molecular pathways were derived using pathway analysis (Figure 2) with GO (Supplementary Table 2) and MP terms (Supplementary Table 3). The resulting 189 GO terms and 118 MP terms were refined to identify terms independently overrepresented among FMRP targets. This procedure left a total of 35 independent overrepresented terms (Supplementary Table 4).

**Figure 2.**
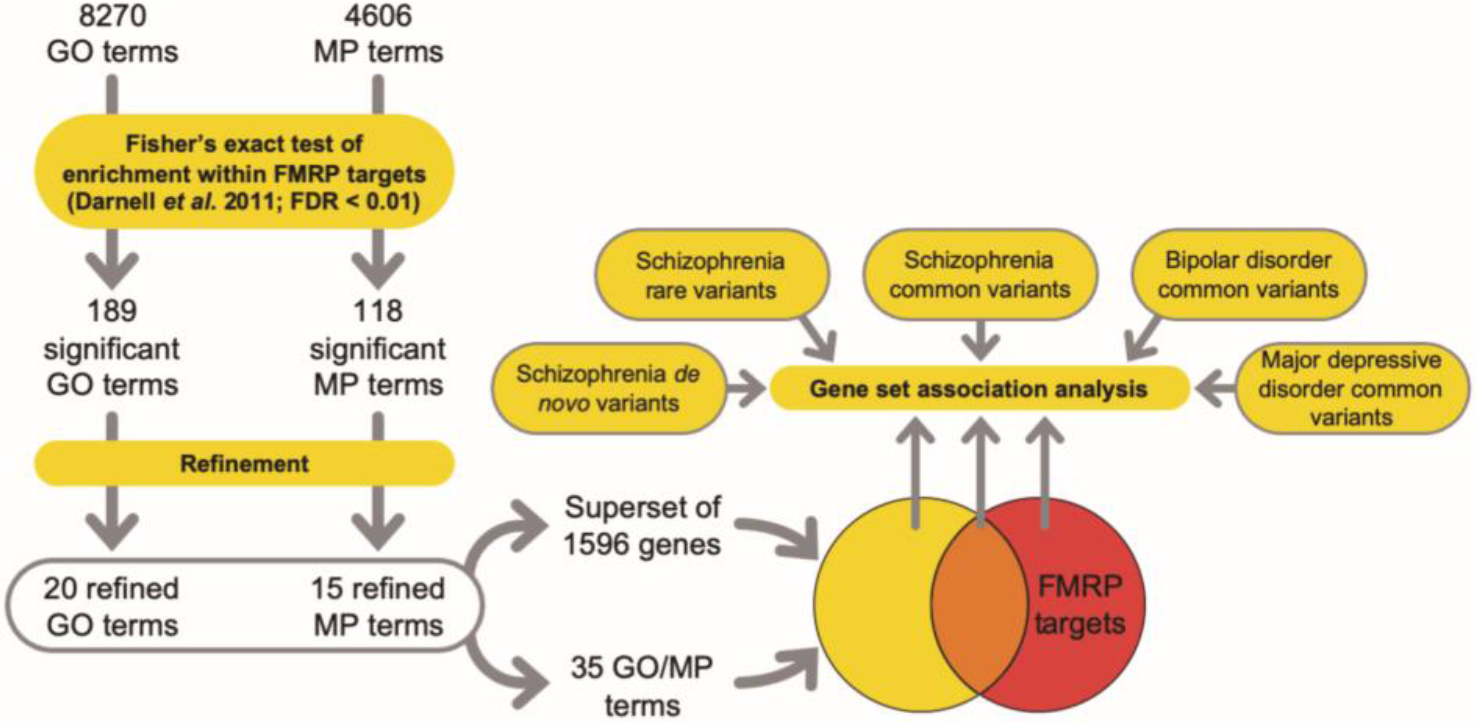
Pathway analysis workflow. Predominant functional subsets of FMRP targets were tested for genetic association with psychiatric disorders. GO, gene ontology; MP, mammalian phenotype; FDR, false discovery rate.

To assess the contribution to genetic association of the property “FMRP binding”, versus that of these functional ontologies, we created a superset (N = 1596) of brain-expressed genes which are included in at least one of the 35 functional terms overrepresented for FMRP targets. FMRP targets from this set (N = 401) were strongly enriched for common variant association (β = 0.29, corrected *P* = 3.7 × 10^−6^), whilst genes not targeted by FMRP (N = 1195) were not (β = 0.066, corrected *P* = 0.13) (Table 1). FMRP targets that were not included in any of the 35 terms (N = 438) were also significantly associated (β = 0.17, corrected *P* = 0.0063). Thus, FMRP targets appear to capture schizophrenia associated genes from these functional categories (when taken as a whole). The burden of rare loss-of-function variants in cases showed the same pattern of association as the common variants, being only enriched in the sets that included FMRP targets (Table 1), regardless of superset membership. However, enrichment for *de novo* nonsynonymous mutations showed a different picture, with significant association being observed only for the set of genes that were exclusive to FMRP targets (Rate ratio = 1.58, corrected *P* = 9.2 × 10^−4^).

**Table 1.** Partitioning FMRP targets genetic association by overrepresented functional annotation. GO and MP functional terms independently overrepresented among FMRP targets were merged, then divided by FMRP targets membership. Genes not brain-expressed were removed. Background association originating from brain expression was controlled for within gene set association analyses. Shown are the resulting effect sizes (β or Rate Ratio) and *P*-values (P). For each variant type, *P*-values were Bonferroni adjusted for 3 tests. SNPs, single nucleotide polymorphisms; LoF, loss-of-function; NS, nonsynonymous.

| Gene set | *N* | Common SNPs |  | Rare LoF |  | *De novo* NS |  |
| --- | --- | --- | --- | --- | --- | --- | --- |
| | | $\beta$ | *P* | $\beta$ | *P* | Rate Ratio | *P* |
| Genes exclusive to functional terms | 1195 | 0.066 | 0.13 | 0.010 | 1.0 | 0.98 | 1.0 |
| Overlapping genes | 401 | 0.29 | $3.7 \times 10^{-6}$ | 0.43 | $3.5 \times 10^{-5}$ | 1.27 | 0.17 |
| Genes exclusive to FMRP targets | 438 | 0.17 | 0.0063 | 0.34 | 0.0023 | 1.58 | $9.2 \times 10^{-4}$ |

In comparisons of effect sizes from analyses of any type of genetic variant, FMRP targets annotated by overrepresented functional terms were not more enriched for association with schizophrenia than unannotated FMRP targets (common variants: *P* = 0.081; rare loss-of-function variants: *P* = 0.25; *de novo* nonsynonymous variants: *P* = 0.88).

We next sought to determine from which of the individual overrepresented functional terms FMRP targets capture genetic association with schizophrenia, and whether association is further enriched within FMRP targets from any single overrepresented term, compared to the complete FMRP targets set. Several functionally-defined subsets of FMRP targets were significantly associated with schizophrenia through common variation (Table 2), whilst genes not targeted by FMRP were not associated; with the exception of those belonging to the term, *calcium ion transmembrane transporter activity* (Supplementary Table 4), although in this instance the fraction targeted by FMRP was associated with a significantly greater effect size (*P* = 0.0088). The *calcium ion transmembrane transporter activity* fraction of FMRP targets (N = 25) remained significantly associated with schizophrenia even after conditioning on all FMRP targets (Supplementary Table 4), implying that this functionally-defined subset of FMRP targets is more enriched for association with schizophrenia than FMRP targets as a whole. No other term captured FMRP targets with a significantly greater enrichment of genetic association than the full FMRP targets gene set.

**Table 2.** GO and MP terms overrepresented among FMRP targets which capture a significant (Padj < 0.05) portion of the common variant genetic association with schizophrenia. Shown are effect sizes (Beta) and *P*-values (P) in gene set association analysis of genes targeted, or not targeted, by FMRP. *P*-values were Bonferroni adjusted (Padj) for 35 terms.

| Term | Genes not FMRP targets |  |  |  | Genes FMRP targets |  |  |  |
| --- | --- | --- | --- | --- | --- | --- | --- | --- |
|  | N | Beta | P | Padj | N | Beta | P | Padj |
| Calcium ion transmembrane transporter activity (GO:0015085) | 91 | 0.419 | $4.7 \times 10^{-4}$ | **0.017** | 25 | 1.080 | $6.9 \times 10^{-6}$ | **$2.4 \times 10^{-4}$** |
| Abnormal motor coordination/balance (MP:0001516) | 538 | 0.104 | 0.028 | 0.97 | 117 | 0.463 | $2.8 \times 10^{-5}$ | **$9.6 \times 10^{-4}$** |
| Abnormal seizure response to inducing agent (MP:0009357) | 125 | 0.190 | 0.043 | 1.0 | 42 | 0.710 | $1.1 \times 10^{-4}$ | **0.0038** |
| Abnormal spatial learning (MP:0001463) | 141 | 0.161 | 0.057 | 1.0 | 61 | 0.569 | $1.4 \times 10^{-4}$ | **0.0049** |
| Growth cone (GO:0030426) | 50 | 0.245 | 0.077 | 1.0 | 27 | 0.854 | $1.7 \times 10^{-4}$ | **0.0060** |
| Abnormal nest building behaviour (MP:0001447) | 15 | 0.265 | 0.22 | 1.0 | 12 | 1.290 | $2.0 \times 10^{-4}$ | **0.0071** |
| Abnormal excitatory postsynaptic currents (MP:0002910) | 60 | 0.177 | 0.12 | 1.0 | 35 | 0.715 | $3.5 \times 10^{-4}$ | **0.012** |
| Axon part (GO:0033267) | 108 | 0.134 | 0.12 | 1.0 | 54 | 0.505 | $6.6 \times 10^{-4}$ | **0.023** |

Rare loss-of-function variants from patients with schizophrenia were enriched in FMRP targets from two terms (*abnormal spatial learning, abnormal motor coordination/balance*) (Supplementary Table 5), whilst no association was found between rare coding variants in non-targeted genes from each term and schizophrenia. None of these subsets harboured significantly more enrichment for case variants than all FMRP targets.

None of the subsets tested captured a significant burden of case *de novo* nonsynonymous variants (Supplementary Table 6).

Overall, these analyses suggest that the overrepresentation of FMRP targets drives genetic association of these biological pathways with schizophrenia, rather than the reverse.

### Genetic association of FMRP targets in other psychiatric disorders

Schizophrenia shares substantial genetic susceptibility with bipolar disorder and major depressive disorder (31–34) and FMRP targets have been previously associated through common variation with major depressive disorder (21). For comparison across disorders, we tested the enrichment of FMRP targets bins for association with major depressive disorder and bipolar disorder using common variant data from GWAS. In both sets of analyses, there was a clear relationship between FMRP binding confidence and genetic association (Figure 3b,c). The topmost bin, containing genes most likely to be FMRP targets, was the most strongly enriched for association with bipolar disorder (corrected *P* = 1.4 × 10^−6^) and major depressive disorder (corrected *P* = 2.5 × 10^−4^).

**Figure 3.**
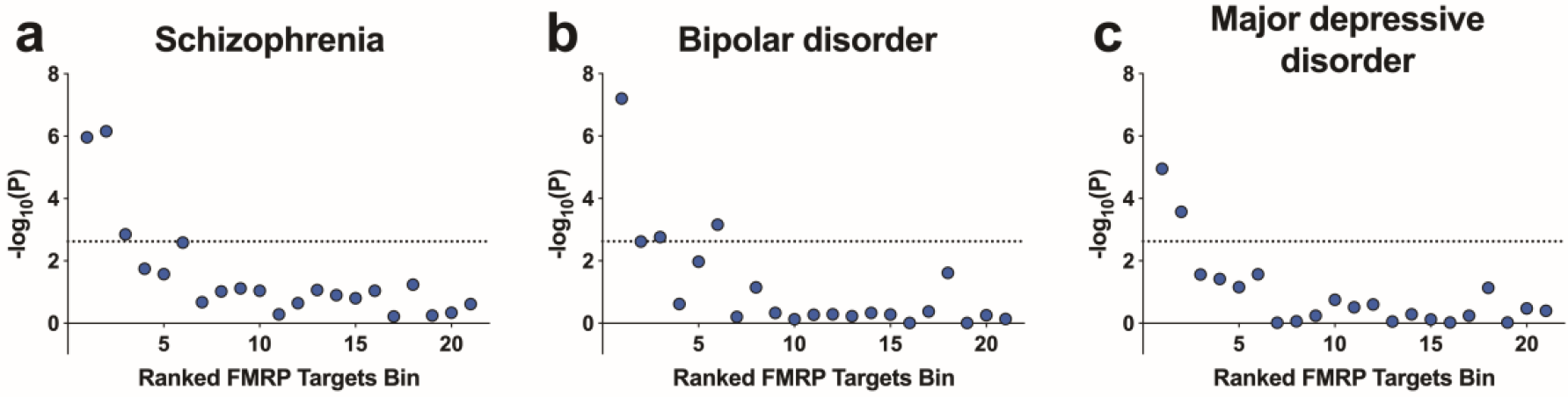
Genetic association of FMRP target bins with schizophrenia, bipolar disorder and major depressive disorder. Shown are −log_10_(*P*-value) following common variant gene set association analysis of 21 bins of 400 genes ranked by FMRP binding confidence. Dotted lines represent a threshold for statistical significance after correction for 21 tests.

We investigated functionally-annotated subgroups of FMRP targets for association with bipolar disorder and major depressive disorder. Beyond background association from brain-expressed genes, FMRP targets annotated for membership of the 35 overrepresented pathways were strongly associated with bipolar disorder (β = 0.23, corrected *P* = 1.6 × 10^−5^) and major depressive disorder (β = 0.21, corrected *P* = 1.6 × 10^−5^), whilst genes from the same functional terms not targeted by FMRP harboured no significant association (bipolar disorder: β = 0.037, corrected *P* = 0.38; major depressive disorder: β = 0.031, corrected *P* = 0.49)(Table 3). A similar picture was observed for individual overrepresented GO / MP terms. Following multiple testing correction, FMRP targets were significantly associated with bipolar disorder from 4 terms (*calcium ion transmembrane transporter activity, abnormal nest building behavior, abnormal spatial learning* and *abnormal seizure response to inducing agent*). Notably, the association of FMRP targets from these 4 terms was common to schizophrenia and bipolar disorder. FMRP targets from 1 term (*abnormal synaptic vesicle morphology*) were significantly associated with major depressive disorder (Supplementary Table 4). FMRP targets belonging to the term *abnormal nest building behavior* (N = 12) were more highly enriched for association with bipolar disorder than FMRP targets as a whole. No FMRP targets were significantly more enriched for association with major depressive disorder than the full FMRP targets set (Supplementary Table 4).

**Table 3.**
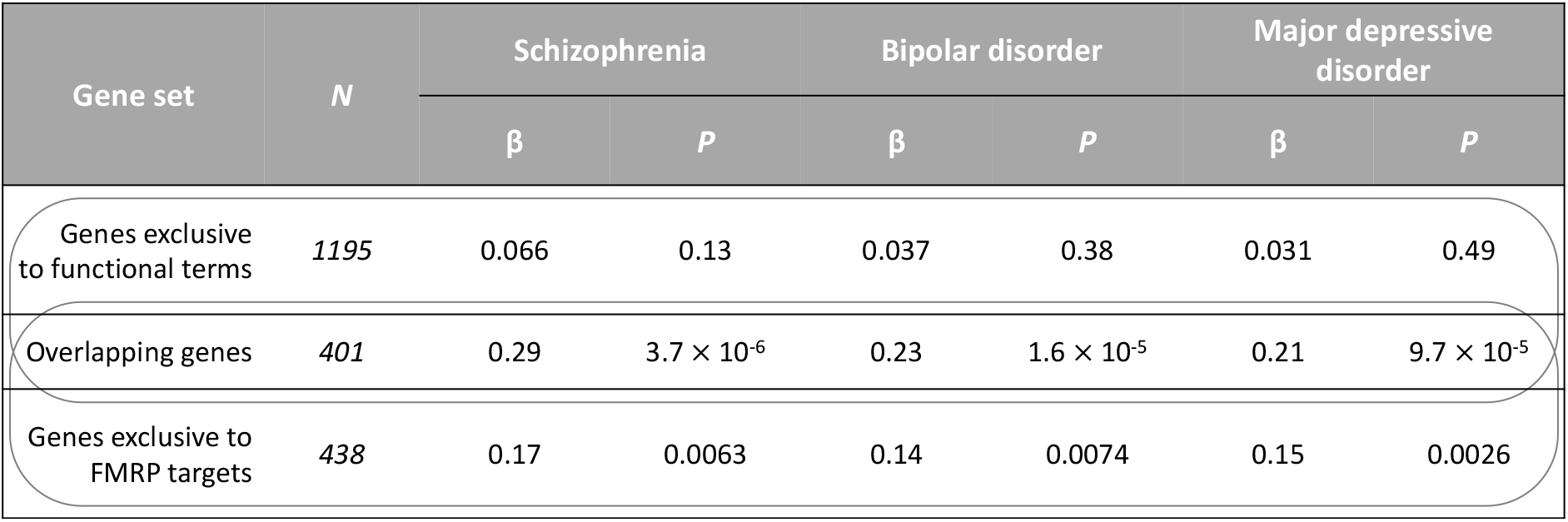
Partitioning FMRP targets common variant association by overrepresented functional annotation. Analyses were performed using a background of brain-expressed genes to account for background association. Shown are the effect sizes (β) and *P*-values (P) from gene set association analyses using MAGMA. For each disorder, *P*-values were adjusted for 3 genes sets using the Bonferroni method.

| Gene set | *N* | Schizophrenia |  | Bipolar disorder |  | Major depressive disorder |  |
| --- | --- | --- | --- | --- | --- | --- | --- |
| | | $\beta$ | *P* | $\beta$ | *P* | $\beta$ | *P* |
| Genes exclusive to functional terms | 1195 | 0.066 | 0.13 | 0.037 | 0.38 | 0.031 | 0.49 |
| Overlapping genes | 401 | 0.29 | $3.7 \times 10^{-6}$ | 0.23 | $1.6 \times 10^{-5}$ | 0.21 | $9.7 \times 10^{-5}$ |
| Genes exclusive to FMRP targets | 438 | 0.17 | 0.0063 | 0.14 | 0.0074 | 0.15 | 0.0026 |

## Discussion

In this study we investigated the extent to which targeting by FMRP is related to genetic association with psychiatric disorders. We show that genes with high probability of being targets of FMRP are enriched for association with schizophrenia, bipolar disorder and major depressive disorder. We also show that it is the property of being an FMRP target that captures the genetic association, rather than membership of gene sets that happen to be enriched for targets of FMRP.

Only bins of genes with the highest FMRP binding confidence were enriched for association with schizophrenia through common variation, exome sequencing-derived rare variation and exome sequencing-derived *de novo* rare variation. This same relationship was reflected in analyses of bipolar disorder and major depressive disorder risk alleles. Our observations are consistent with previous gene set analyses of FMRP targets in the context of schizophrenia (11–15) and major depressive disorder (21), but whilst FMRP targets have been previously linked to bipolar disorder through rare coding variants (35), our findings provide novel evidence linking FMRP targets to bipolar disorder through common variation.

Despite the evidence implicating FMRP targets in psychiatric disorders (11–15), the overrepresentation of long, brain-expressed genes with synaptic functions has led to cautiousness over the validity of the link to FMRP (22). The methods used here, and previously (11), correct for, or are unaffected by, gene length, allowing us more confidence in concluding that the relationships between FMRP binding and association with schizophrenia, bipolar disorder and major depressive disorder exist beyond any confounding effects of gene length. Furthermore, whilst the associated genes were derived from expressed mRNAs in mouse brain, the associations did not generalize to bins of brain-expressed genes with low FMRP binding confidence.

Consistent with previous pathway analysis (1), we note that a substantial proportion of FMRP targets have functions related to synaptic activity, anatomy or development. Studies of FMRP function show that its activity is regulated in response to neuronal activity (36–39) and is an important mediator of synapse development (40–42), synaptic plasticity (43–45), learning and memory (46–48). Genetic and functional studies have highlighted the relevance of perturbed synaptic plasticity in psychiatric disorders (12,30,49–53), although we find that the risk conferred by variants affecting such pathways overrepresented among FMRP targets is concentrated within the fraction of genes targeted by FMRP. Hence, despite the convergence of psychiatric risk on synaptic pathways (12,23,51–54), the association of FMRP targets was not attributed to these overrepresented annotations. Instead, it appears that there is a degree of specificity to this risk, such that genes regulated locally by FMRP during activity-induced synaptic plasticity, required for development or learning, are most relevant to psychiatric disorder.

It should be noted that other synapse-related gene sets are enriched for association with psychiatric disorders independently of FMRP targets. For instance, recent schizophrenia common variant analyses show a few such independent associations of gene sets related to synaptic function (11). Here we found that, whilst strongest for genes targeted by FMRP, genes involved in *calcium ion transmembrane transporter activity* held independent association with schizophrenia. Furthermore, the strongly associated, albeit small, intersection between genes from this set and FMRP targets contained a stronger enrichment of schizophrenia common variant association than FMRP targets (or indeed the GO term) as a whole. This is consistent with previous evidence for association of calcium channels with schizophrenia (10,11,13), yet additionally suggests that FMRP captures a subset of genes related to calcium ion transport in which common variant association is concentrated.

FMRP binding confidence was not related to genetic association with schizophrenia through CNVs. Whilst FMRP targets have been consistently implicated in schizophrenia from analyses of all other types of genetic variant, studies of structural variation in schizophrenia have shown only modest association of FMRP targets (28,30,55), although a deletion at 15q11.2 affecting the FMRP interacting protein, CYFIP1, which is required for the regulation of translation by FMRP (5,56), is associated. Why we observe risk for schizophrenia affecting FMRP targets being conferred through all variants except for CNVs is unclear, although these analyses may be influenced by the difficulty of attributing CNV association to individual genes.

Our observations resonate with the growing body of literature challenging the biological validity of viewing major psychiatric disorders as discrete entities with independent genetic aetiology (57–60). There is considerable overlap between the genetic risk attributable to schizophrenia, bipolar disorder and major depressive disorder (31–34). The present (and published) findings highlight that FMRP targets are a point of shared heritability. Additional evidence suggests that genetic association of FMRP targets may extend also to autism (17–20) and attention-deficit hyperactivity disorder (61).

Our findings highlight a set of genes regulated through a common mechanism that harbour risk across several psychiatric disorders. However, there is a degree of uncertainty as to precisely which mRNAs are regulated by FMRP. Multiple studies have examined this, each yielding overlapping, yet distinct sets of FMRP targets (1,4,62–65); some of the variability likely originating from tissue-specificity. When performing pathway analyses with genomic data, many studies, including this one, have obtained FMRP targets from an investigation of mRNA-FMRP interaction sites in mouse cortical polyribosomes (1), in which membership was assigned by applying a stringent cut-off to a continuous scale of binding confidence, likely resulting in some false positives and more false negatives. Moreover, binding by FMRP may not equate to translational repression in the cell, which requires additional contribution from binding partners CYFIP1 and eIF4E, within a protein complex (5). Hence, this line of research will benefit from further validation of FMRP-regulated protein synthesis in the context of psychiatric pathology.

Our results serve to strengthen the evidence that a population of genes targeted by FMRP, many of which have synaptic functions, are affected by genetic variation conferring risk to psychiatric disorders, including schizophrenia, bipolar disorder and major depressive disorder. We conclude that targeting by FMRP is currently the most suitable functional annotation to reflect the origin of these associations and represents a common mode of regulation for a set of genes contributing risk across several major psychiatric presentations.

## Materials and Methods

### Gene sets

FMRP binding statistics for 30999 transcripts were obtained from Darnell *et al* (2011) (Supplementary Table S2C) (1), a study of mRNA-FMRP interaction sites in mouse cortical polyribosomes using crosslinking immunoprecipitation combined with high-throughput RNA sequencing. We filtered the data to include only genes which were expressed (chi-square score > 0), therefore selecting only those from which binding statistics could be obtained. We converted Mouse Entrez IDs to human Entrez IDs *via* their shared HomoloGene ID, obtained from Mouse Genome Informatics Vertebrate Homology database release 6.10 (HOM_AllOrganism.rpt, 8^th^ January 2018). Genes that did not convert to a unique protein coding human homologue were excluded. The remaining 8595 genes were ranked by their FMRP binding confidence *P*-value and the top 8400 were split into 21 bins of 400 genes to determine the relationship between FMRP binding confidence and schizophrenia association. Functional enrichment analyses were performed using the set of 842 FMRP targets (reported FDR < 0.01 in Darnell *et al*, 2011) (1) that has been widely used in previous enrichment studies (11,12).

### samples

#### Common variants

All genetic data were obtained from published case and control samples. Schizophrenia common variant summary statistics were taken from the Pardiñas *et al* (2018) study, a meta analysis of genome-wide association studies (GWAS) (11) based on a sample of 40 675 case and 64 643 control subjects. Bipolar disorder common variant summary statistics were provided by a recent Psychiatric Genomics Consortium (PGC) GWAS (53), consisting of 20 352 cases and 31 358 controls from 32 cohorts of European descent. Major depressive disorder common variant summary statistics were taken from a PGC meta-analysis of 135 458 cases and 344 901 controls from seven independent cohorts of European ancestry (21).

#### Rare coding variants

Exome sequencing-derived rare coding variant data from a Swedish schizophrenia case control study (16) were obtained from the NCBI database of genotypes and phenotypes (dbGaP). After excluding individuals with non-European or Finnish ancestry, and samples with low sequencing coverage, we retained exome sequence in 4079 cases and 5712 controls for analysis.

#### De novo coding variants

*De novo* mutations were derived (66) from previously published exome sequencing studies of, collectively, 1136 schizophrenia-proband parent trios (12,67–74) (Supplementary Table 1).

#### Copy number variants

Copy number variant (CNV) data were compiled from the CLOZUK and Cardiff Cognition in Schizophrenia samples (11 955 cases, 19 089 controls) (27,75), as well as samples from the International Schizophrenia Consortium (3395 cases, 2185 controls) (76) and the Molecular Genetics of Schizophrenia (2215 cases, 2556 controls) (77), giving a total of 17 565 case and 24 830 control subjects. Genotyping, CNV calling and quality control information can be found in the original reports (25,27,30,49,76,77).

### Gene set association analysis

Schizophrenia, bipolar disorder and major depressive disorder GWAS single nucleotide polymorphisms (SNPs) were filtered to include only those with a minor allele frequency ≥ 0.01. SNP association *P*-values were combined (SNP-wise Mean model) into gene-wide *P-* values in MAGMA v1.06 (24), using a window of 35 kb upstream and 10 kb downstream of each gene to include proximal regulatory regions. The European panel of the 1000 Genomes Project (78) (phase 3) was used as a reference to account for linkage disequilibrium between genes. Gene sets were tested for enrichment for association with each disorder using one-tailed competitive gene set association analyses in MAGMA, which compares the mean association of genes from the gene set to those not in the gene set, correcting for gene size and SNP density. The default background was all protein-coding genes.

Case-control exome sequencing data were analysed using Hail (https://github.com/hail-is/hail). We annotated variants using Hail’s Ensembl VEP method (version 86, http://oct2016.archive.ensembl.org/index.html) and defined loss-of-function variants as nonsense, essential splice site and frameshift annotations and nonsynonymous variants as loss-of-function and missense annotations. For gene set enrichment tests, we focused on ultra-rare singleton loss-of-function and nonsynonymous variants, that is those observed once in all case-control sequencing data and absent from the non-psychiatric component of ExAC (79). Enrichment statistics were generated using a Firth’s penalized-likelihood logistic regression model that corrected for the first 10 principal components, exome-wide burden of synonymous variants, sequencing platform and sex.

*De novo* variant gene set enrichment was evaluated by comparing the observed number of *de novo* variants in a set of genes to that expected, which was based on the number of trios analysed and per-gene mutations rates (80,81). Gene set enrichment statistics for *de novo* variants were generated by using a two sample Poisson rate ratio test to compare the enrichment of *de novo* variants within the gene set to that observed in a background set of genes.

CNV analyses were restricted to CNVs at least 100 kb in size and covered by at least 15 probes. Gene set association was tested by logistic regression, in which CNV case-control status was regressed against the number of set genes overlapped by the CNV, with covariates: CNV size, genes per CNV, study and chip type. To correct for *P*-value inflation, empirical *P*-values were obtained by calculating the fraction of random size-matched sets of brain-expressed (1) genes that yielded an association as or more significant.

Multiple testing was corrected for using the Bonferroni method.

### Pathway analysis

For gene ontology enrichment analyses, functional annotations of each gene were compiled separately from the Gene Ontology (GO) (82) and Mouse Genome Informatics (MGI) Mammalian Phenotype (MP) (83) databases (July 4^th^ 2018). GO annotations were filtered to exclude genes with the following evidence codes: NAS (Non-traceable Author Statement), IEA (Inferred from Electronic Annotation), and RCA (inferred from Reviewed Computational Analysis). GO or MP terms containing fewer than 10 genes were then excluded. For all pathway analyses, genes were restricted to those expressed (chi-square score > 0) in the mouse brain tissue used by Darnell *et al* 2011 (1). Enrichment of FMRP targets for each GO/MP term was assessed by Fisher’s exact tests, with the contrast group being all remaining brain-expressed genes. Following separate Bonferroni correction for 8270 GO terms or 4606 MP terms, significantly (*P* < 0.01) overrepresented terms were subjected to a competitive refinement procedure to resolve the effects of redundancy between terms. During refinement, terms were re-tested for overrepresentation in FMRP targets following the removal of genes from the term with the highest odds ratio in Fisher’s exact test. Terms that were no longer significant upon re-test (unadjusted *P* > 0.01) were dropped. This was done repeatedly, such that genes from the remaining term with the highest odds ratio on each repeat were removed in addition to those removed on previous iterations.

In primary analyses of genetic association, brain-expressed (1) genes from all overrepresented GO / MP terms (following refinement) were grouped together and divided into those targeted and those not targeted by FMRP, and compared to a background of brain-expressed genes. In secondary analyses, genes from each individual overrepresented term were divided in the same way and tested for association using all protein-coding genes as a comparator. *P*-values were Bonferroni corrected for the number of functional terms being tested at each stage of analysis.

We performed a number of tests to investigate the relative enrichments for association between two sets of genes, one a subset of the other. For common variant association, we used the conditional analysis function provided by MAGMA. For rare or *de novo* coding variants, we compared the effect sizes of the subset of genes with that of the larger set after excluding members of the subset. For the rare coding variant case control analyses, this was done by performing a z-test of beta values, whilst for *de novo* variant analyses, a two-sample Poisson rate ratio test was used.

In cases where enrichment for genetic association was compared between non-overlapping gene sets, a z-test of beta values (common and rare variants) or a two-sample Poisson rate ratio test (*de novo* variants) was used.

## Acknowledgements

This work was supported by Medical Research Council (MRC) grants MR/L010305/1 and G0800509, a Wellcome Trust Strategic Award (100202/Z/12/Z), The Waterloo Foundation ‘Changing Minds’ programme, and Neuroscience and Mental Health Research Institute (Cardiff University) core funding to NC.

We thank the Bipolar Disorder and Major Depressive Disorder workgroups of the Psychiatric Genomics Consortium for providing summary statistics used in this study. We would also like to thank the research participants and employees of 23andMe for making this work possible. Exome sequencing datasets described in this manuscript were obtained from dbGaP at http://www.ncbi.nlm.nih.gov/gap through dbGaP accession number phs000473.v2.p2. Samples were provided by the Swedish Cohort Collection supported by the NIMH Grant No. R01MH077139, the Sylvan C. Herman Foundation, the Stanley Medical Research Institute and The Swedish Research Council (Grant Nos. 2009-4959 and 2011-4659). Support for the exome sequencing was provided by the NIMH Grand Opportunity Grant No. RCMH089905, the Sylvan C. Herman Foundation, a grant from the Stanley Medical Research Institute and multiple gifts to the Stanley Center for Psychiatric Research at the Broad Institute of MIT and Harvard.

Analyses of copy number variation described in this manuscript used datasets from the Molecular Genetics of Schizophrenia (MGS; dbGAP phs000021.v3.p2 and phs000167.v1.p1) and the International Schizophrenia Consortium (ISC). The CLOZUK and CLOZUK2 data sets contain data obtained from outside sources: dbGaP phs000404.v1.p1, phs000187.v1. p1, phs000303.v1.p1, phs000179.v3.p2, phs000421.v1.p, phs000395.v1.p1, phs000519.v1.p1 and the Wellcome Trust Case Control Consortium 2 study.

## Abbreviations

FMRP: Fragile X mental retardation protein
mRNA: Messenger ribonucleic acid
MAGMA: Multi-marker Analysis of GenoMic Annotation
GWAS: Genome-wide association study
DNA: Deoxyribonucleic acid
CNV: Copy number variant
GO: Gene ontology
MGI: Mouse genome informatics
MP: Mammalian phenotype
CYFIP1: Cytoplasmic FMR1 interacting protein 1
PGC: Psychiatric genomics consortium

## References

1. Darnell, J. C., Van Driesche, S. J., Zhang, C., et al. (2011) FMRP stalls ribosomal translocation on mRNAs linked to synaptic function and autism. Cell, 146, 247–261.

2. Laggerbauer, B., Ostareck, D., Keidel, E. M., et al. (2001) Evidence that fragile X mental retardation protein is a negative regulator of translation. Hum. Mol. Genet.

3. Li, Z., Zhang, Y., Ku, L., et al. (2001) The fragile X mental retardation protein inhibits translation via interacting with mRNA. Nucleic Acids Res.

4. Ascano, M., Mukherjee, N., Bandaru, P., et al. (2012) FMRP targets distinct mRNA sequence elements to regulate protein expression. Nature, 492, 382–6.

5. Napoli, I., Mercaldo, V., Boyl, P. P., et al. (2008) The Fragile X Syndrome Protein Represses Activity-Dependent Translation through CYFIP1, a New 4E-BP. Cell, 134, 1042–1054.

6. Christie, S. B., Akins, M. R., Schwob, J. E., et al. (2009) The FXG: A Presynaptic Fragile X Granule Expressed in a Subset of Developing Brain Circuits. J. Neurosci.

7. Stefani, G., Fraser, C. E., Darnell, J. C., et al. (2004) Fragile X Mental Retardation Protein Is Associated with Translating Polyribosomes in Neuronal Cells. J. Neurosci.

8. Verkerk, A. J. M. H., Pieretti, M., Sutcliffe, J. S., et al. (1991) Identification of a gene (FMR-1) containing a CGG repeat coincident with a breakpoint cluster region exhibiting length variation in fragile X syndrome. Cell, 65, 905–914.

9. Bagni, C., Tassone, F., Neri, G., et al. (2012) Fragile X syndrome: Causes, diagnosis, mechanisms, and therapeutics. Fragile X syndrome: Causes, diagnosis, mechanisms, and therapeutics. J. Clin. Invest. (2012).

10. Ripke, S., Neale, B. M., Corvin, A., et al. (2014) Biological insights from 108 schizophrenia-associated genetic loci. Nature, 511, 421–427.

11. Pardiñas, A. F., Holmans, P., Pocklington, A. J., et al. (2018) Common schizophrenia alleles are enriched in mutation-intolerant genes and in regions under strong background selection. Nat. Genet., 50, 381–389.

12. Fromer, M., Pocklington, A. J., Kavanagh, D. H., et al. (2014) De novo mutations in schizophrenia implicate synaptic networks. Nature, 506, 179–84.

13. Purcell, S. M., Moran, J. L., Fromer, M., et al. (2014) A polygenic burden of rare disruptive mutations in schizophrenia. Nature, 506.

14. Richards, A. L., Leonenko, G., Walters, J. T., et al. (2016) Exome arrays capture polygenic rare variant contributions to schizophrenia. Hum. Mol. Genet., 25, 1001–1007.

15. Leonenko, G., Richards, A. L., Walters, J. T., et al. (2017) Mutation intolerant genes and targets of FMRP are enriched for nonsynonymous alleles in schizophrenia. Am. J. Med. Genet. Part B Neuropsychiatr. Genet., 724–731.

16. Genovese, G., Fromer, M., Stahl, E. A., et al. (2016) Increased burden of ultra-rare protein altering variants among 4,877 individuals with schizophrenia. Nat. Neurosci., 19, 1433–1441.

17. Jansen, A., Dieleman, G. C., Smit, A. B., et al. (2017) Gene-set analysis shows association between FMRP targets and autism spectrum disorder. Eur. J. Hum. Genet., 25, 863–868.

18. Iossifov, I., Ronemus, M., Levy, D., et al. (2012) De Novo Gene Disruptions in Children on the Autistic Spectrum. Neuron.

19. Iossifov, I., O’Roak, B. J., Sanders, S. J., et al. (2014) The contribution of de novo coding mutations to autism spectrum disorder. Nature.

20. De Rubeis, S., He, X., Goldberg, A. P., et al. (2014) Synaptic, transcriptional and chromatin genes disrupted in autism. Nature.

21. Wray, N., Ripke, S., Mattheisen, M., et al. (2018) Genome-wide association analyses identify 44 risk variants and refine the genetic architecture of major depression. Nat. Genet., 50, 668–681.

22. Ouwenga, R. L. and Dougherty, J. (2015) Fmrp targets or not: long, highly brain-expressed genes tend to be implicated in autism and brain disorders. Mol. Autism, 6, 16.

23. Hall, J., Trent, S., Thomas, K. L., et al. (2015) Genetic risk for schizophrenia: Convergence on synaptic pathways involved in plasticity. Biol. Psychiatry, 77, 52–58.

24. de Leeuw, C. A., Mooij, J. M., Heskes, T., et al. (2015) MAGMA: Generalized Gene-Set Analysis of GWAS Data. PLoS Comput. Biol., 11.

25. Rees, E., Walters, J. T. R., Georgieva, L., et al. (2014) Analysis of copy number variations at 15 schizophrenia-associated loci. Br. J. Psychiatry, 204, 108–114.

26. Kirov, G., Grozeva, D., Norton, N., et al. (2009) Support for the involvement of large copy number variants in the pathogenesis of schizophrenia. Hum. Mol. Genet., 18, 1497–1503.

27. Rees, E., Walters, J. T. R., Chambert, K. D., et al. (2014) CNV analysis in a large schizophrenia sample implicates deletions at 16p12.1 and SLC1A1 and duplications at 1p36.33 and CGNL1. Hum. Mol. Genet., 23, 1669–1676.

28. Marshall, C. R., Howrigan, D. P., Merico, D., et al. (2017) Contribution of copy number variants to schizophrenia from a genome-wide study of 41,321 subjects. Nat. Genet., 49, 27–35.

29. Kirov, G., Pocklington, A. J., Holmans, P., et al. (2012) De novo CNV analysis implicates specific abnormalities of postsynaptic signalling complexes in the pathogenesis of schizophrenia. Mol. Psychiatry, 17, 142–153.

30. Pocklington, A. J., Rees, E., Walters, J. T. R., et al. (2015) Novel Findings from CNVs Implicate Inhibitory and Excitatory Signaling Complexes in Schizophrenia. Neuron, 86, 1203–1214.

31. Lee, S. H., Ripke, S., Neale, B. M., et al. (2013) Genetic relationship between five psychiatric disorders estimated from genome-wide SNPs. Nat. Genet., 45, 984–94.

32. Cardno, A. G. and Owen, M. J. (2014) Genetic relationships between schizophrenia, bipolar disorder, and schizoaffective disorder. Schizophr. Bull., 40, 504–515.

33. Owen, M. J. and O’Donovan, M. C. (2017) Schizophrenia and the neurodevelopmental continuum:evidence from genomics. World Psychiatry, 16, 227–235.

34. Doherty, J. L. and Owen, M. J. (2014) Genomic insights into the overlap between psychiatric disorders: implications for research and clinical practice. Genome Med., 6, 29.

35. Goes, F. S., Pirooznia, M., Parla, J. S., et al. (2016) Exome sequencing of familial bipolar disorder. JAMA Psychiatry, 73, 590–597.

36. Nakamoto, M., Nalavadi, V., Epstein, M. P., et al. (2007) Fragile X mental retardation protein deficiency leads to excessive mGluR5-dependent internalization of AMPA receptors. Proc. Natl. Acad. Sci.

37. Wang, H., Wu, L. J., Kim, S. S., et al. (2008) FMRP Acts as a Key Messenger for Dopamine Modulation in the Forebrain. Neuron.

38. Wang, H., Morishita, Y., Miura, D., et al. (2012) Roles of CREB in the regulation of FMRP by group i metabotropic glutamate receptors in cingulate cortex. Roles of CREB in the regulation of FMRP by group i metabotropic glutamate receptors in cingulate cortex. Mol. Brain (2012).

39. Antar, L. N., Afroz, R., Dictenberg, J. B., et al. (2004) Metabotropic Glutamate Receptor Activation Regulates Fragile X Mental Retardation Protein and Fmr1 mRNA Localization Differentially in Dendrites and at Synapses. J. Neurosci., 24, 2648–2655.

40. Wang, X., Zorio, D. A. R., Schecterson, L., et al. (2018) Postsynaptic FMRP Regulates Synaptogenesis In Vivo in the Developing Cochlear Nucleus. J. Neurosci.

41. Davis, J. K. and Broadie, K. (2017) Multifarious Functions of the Fragile X Mental Retardation Protein. Multifarious Functions of the Fragile X Mental Retardation Protein. Trends Genet. (2017).

42. Doll, C. A. and Broadie, K. (2015) Activity-dependent FMRP requirements in development of the neural circuitry of learning and memory. Development.

43. Shang, Y., Wang, H., Mercaldo, V., et al. (2009) Fragile X mental retardation protein is required for chemically-induced long-term potentiation of the hippocampus in adult mice. J. Neurochem.

44. Huber, K. M., Gallagher, S. M., Warren, S. T., et al. (2002) Altered synaptic plasticity in a mouse model of fragile X mental retardation. Proc. Natl. Acad. Sci. U. S. A., 99, 7746–7750.

45. Eadie, B. D., Cushman, J., Kannangara, T. S., et al. (2012) NMDA receptor hypofunction in the dentate gyrus and impaired context discrimination in adult Fmr1 knockout mice. Hippocampus.

46. Yan, Q. J., Asafo-Adjei, P. K., Arnold, H. M., et al. (2004) A phenotypic and molecular characterization of the fmr1-tm1Cgr fragile X mouse. Genes, Brain Behav.

47. Michalon, A., Sidorov, M., Ballard, T. M., et al. (2012) Chronic Pharmacological mGlu5 Inhibition Corrects Fragile X in Adult Mice. Neuron.

48. Santos, A. R., Kanellopoulos, A. K. and Bagni, C. (2014) Learning and behavioral deficits associated with the absence of the fragile X mental retardation protein: What a fly and mouse model can teach us. Learning and behavioral deficits associated with the absence of the fragile X mental retardation protein: What a fly and mouse model can teach us. Learn. Mem. (2014)

49. Clifton, N. E., Pocklington, A. J., Scholz, B., et al. (2017) Schizophrenia copy number variants and associative learning. Mol. Psychiatry, 22, 178–182.

50. Duman, R. S., Aghajanian, G. K., Sanacora, G., et al. (2016) Synaptic plasticity and depression: New insights from stress and rapid-acting antidepressants. Synaptic plasticity and depression: New insights from stress and rapid-acting antidepressants. Nat. Med. (2016).

51. Howard, D. M., Adams, M. J., Shirali, M., et al. (2018) Genome-wide association study of depression phenotypes in UK Biobank identifies variants in excitatory synaptic pathways. Nat. Commun., 9, 1470.

52. Howard, D. M., Adams, M. J., Clarke, T. K., et al. (2019) Genome-wide meta-analysis of depression identifies 102 independent variants and highlights the importance of the prefrontal brain regions. Nat. Neurosci., 22, 343–352.

53. Stahl, E. A., Breen, G., Forstner, A. J., et al. (2019) Genome-wide association study identifies 30 loci associated with bipolar disorder. Nat. Genet., 51, 793–803.

54. Rees, E., O’Donovan, M. C. and Owen, M. J. (2015) Genetics of schizophrenia. Curr. Opin. Behav. Sci., 2, 8–14.

55. Szatkiewicz, J., O’dushlaine, C., Chen, G., et al. (2014) Copy number variation in schizophrenia in Sweden. Mol. Psychiatry, 19, 762–773.

56. DeRubeis, S., Pasciuto, E., Li, K., et al. (2013) CYFIP1 coordinates mRNA translation and cytoskeleton remodeling to ensure proper dendritic Spine formation. Neuron, 79, 1169–1182.

57. O’Donovan, M. C. and Owen, M. J. (2016) The implications of the shared genetics of psychiatric disorders. The implications of the shared genetics of psychiatric disorders. Nat. Med. (2016), 22.

58. Owen, M. J. (2014) New approaches to psychiatric diagnostic classification. New approaches to psychiatric diagnostic classification. Neuron (2014), 84, 564–571.

59. Craddock, N. and Owen, M. J. (2010) The Kraepelinian dichotomy - Going, going… but still not gone. The Kraepelinian dichotomy - Going, going… but still not gone. Br. J. Psychiatry (2010), 196, 92–95.

60. Jablensky, A. (2016) Psychiatric classifications: Validity and utility. World Psychiatry.

61. Thapar, A., Martin, J., Mick, E., et al. (2015) Psychiatric gene discoveries shape evidence on ADHD’s biology. Mol. Psychiatry, 21, 1202–1207.

62. Suhl, J. a, Chopra, P., Anderson, B. R., et al. (2014) Analysis of FMRP mRNA target datasets reveals highly associated mRNAs mediated by G-quadruplex structures formed via clustered WGGA sequences. Hum. Mol. Genet., 23, 1–41.

63. Brown, V., Jin, P., Ceman, S., et al. (2001) Microarray identification of FMRP-associated brain mRNAs and altered mRNA translational profiles in fragile X syndrome. Cell, 107, 477–487.

64. Maurin, T., Lebrigand, K., Castagnola, S., et al. (2018) HITS-CLIP in various brain areas reveals new targets and new modalities of RNA binding by fragile X mental retardation protein. Nucleic Acids Res., 1–12.

65. Miyashiro, K. Y., Beckel-Mitchener, A., Purk, T. P., et al. (2003) RNA cargoes associating with FMRP reveal deficits in cellular functioning in Fmr1 null mice. Neuron, 37, 417–431.

66. Rees, E., Carrera, N., Morgan, J., et al. (2019) Targeted Sequencing of 10,198 Samples Confirms Abnormalities in Neuronal Activity and Implicates Voltage-Gated Sodium Channels in Schizophrenia Pathogenesis. Biol. Psychiatry, 85, 554–562.

67. Ambalavanan, A., Girard, S. L., Ahn, K., et al. (2016) De novo variants in sporadic cases of childhood onset schizophrenia. Eur. J. Hum. Genet., 24, 944–948.

68. Girard, S. L., Gauthier, J., Noreau, A., et al. (2011) Increased exonic de novo mutation rate in individuals with schizophrenia. Nat. Genet., 43, 860–863.

69. Guipponi, M., Santoni, F. A., Setola, V., et al. (2014) Exome sequencing in 53 sporadic cases of schizophrenia identifies 18 putative candidate genes. PLoS One, 9.

70. Gulsuner, S., Walsh, T., Watts, A. C., et al. (2013) Spatial and temporal mapping of de novo mutations in schizophrenia to a fetal prefrontal cortical network. Cell, 154.

71. McCarthy, S. E., Gillis, J., Kramer, M., et al. (2014) De novo mutations in schizophrenia implicate chromatin remodeling and support a genetic overlap with autism and intellectual disability. Mol. Psychiatry, 19, 652–658.

72. Takata, A., Xu, B., Ionita-Laza, I., et al. (2014) Loss-of-Function Variants in Schizophrenia Risk and SETD1A as a Candidate Susceptibility Gene. Neuron, 82, 773–780.

73. Wang, Q., Li, M., Yang, Z., et al. (2015) Increased co-expression of genes harboring the damaging de novo mutations in Chinese schizophrenic patients during prenatal development. Sci. Rep., 5.

74. Xu, B., Ionita-Laza, I., Roos, J. L., et al. (2012) De novo gene mutations highlight patterns of genetic and neural complexity in schizophrenia. Nat. Genet., 44, 1365–1369.

75. Rees, E., Kendall, K., Pardiñas, A. F., et al. (2016) Analysis of Intellectual Disability Copy Number Variants for Association With Schizophrenia. JAMA Psychiatry, 73, 963–969.

76. Stone, J. L., O’Donovan, M. C., Gurling, H., et al. (2008) Rare chromosomal deletions and duplications increase risk of schizophrenia. Nature, 455, 237–241.

77. Levinson, D. F., Duan, J., Oh, S., et al. (2011) Copy number variants in schizophrenia: Confirmation of five previous finding sand new evidence for 3q29 microdeletions and VIPR2 duplications. Am. J. Psychiatry, 168, 302–316.

78. Auton, A., Abecasis, G. R., Altshuler, D. M., et al. (2015) A global reference for human genetic variation. Nature, 526, 68–74.

79. Lek, M., Karczewski, K. J., Minikel, E. V, et al. (2016) Analysis of protein-coding genetic variation in 60,706 humans. Nature.

80. Samocha, K. E., Robinson, E. B., Sanders, S. J., et al. (2014) A framework for the interpretation of de novo mutation in human disease. Nat. Genet., 46, 944–950.

81. Ware, J. S., Samocha, K. E., Homsy, J., et al. (2015) Interpreting de novo Variation in Human Disease Using denovolyzeR. Curr. Protoc. Hum. Genet.

82. The Gene Ontology Consortium (2017) Expansion of the Gene Ontology knowledgebase and resources. Nucleic Acids Res.

83. Smith, C. L., Blake, J. A., Kadin, J. A., et al. (2018) Mouse Genome Database (MGD)-2018: Knowledgebase for the laboratory mouse. Nucleic Acids Res., 46, D836–D842.

