## Supplementary Info for "Genetic association of FMRP targets with psychiatric disorders"

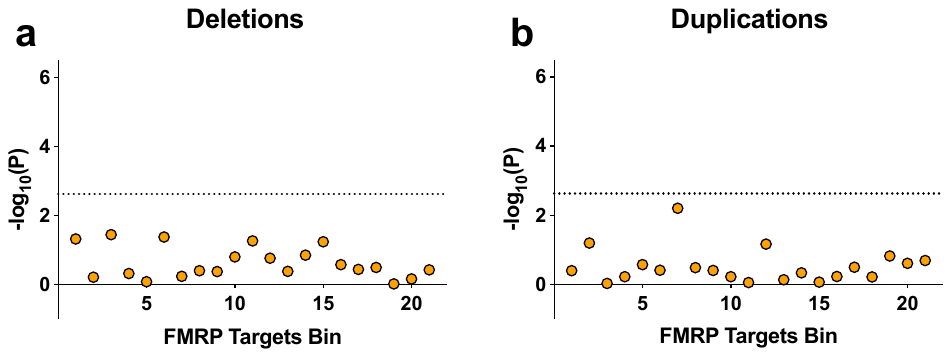

Supplementary Figure 1 Genetic association with schizophrenia of FMRP target bins, following separation of copy number variants (CNVs) into deletions and duplications. Shown are -log_10_(P­-value), where P-values were empirically derived using gene set association analysis, adjusting for inflation observed through parallel analyses of random size-matched sets of expressed genes.

| Study | N probands (Male:Female) |
| --- | --- |
| Fromer *et al* 2014 | 617 (302:315) |
| Girard *et al* 2011 | 14 (7:7) |
| Xu *et al* 2012 / Takata *et al* 2014 | 231 (156:75) |
| Gulsuner *et al* 2013 | 105 (75:30) |
| Wang *et al* 2015 | 45 (22:23) |
| Ambalavanan *et al* 2015 | 17 (11:6) |
| Guipponi *et al* 2014 | 53 (39:14) |
| McCarthy *et al* 2014 | 54 (43:11) |

Supplementary Table 1. Sources of published schizophrenia de novo variants.
